# A Comparative Evaluation of Computational Models for RNA modification detection using Nanopore sequencing with RNA004 Chemistry

**DOI:** 10.1101/2025.02.03.636352

**Authors:** Yongji Zou, Mian Umair Ahsan, Joe Chan, Wen Meng, Shou-Jiang Gao, Yufei Huang, Kai Wang

## Abstract

Direct RNA sequencing from Oxford Nanopore Technologies (ONT) has become a valuable method for studying RNA modifications such as N6-methyladenosine (m6A), pseudouridine (ψ), and 5-methylcytosine (m5C). Recent advancements in the RNA004 chemistry substantially reduce sequencing errors compared to previous chemistries (e.g., RNA002), thereby promising enhanced accuracy for epitranscriptomic analysis. In this study, we benchmark the performance of two state-of-the-art RNA modification detection models capable of handling RNA004 data - ONT’s Dorado and m6Anet - using two wild-type (WT) cell lines, HEK293T and HeLa, with respective ground truths from GLORI and eTAM-seq, and their paired in vitro transcribed (IVT) RNA as negative controls. We found that under default settings and considering sites with ≥10% modification ratio and ≥10X coverage, Dorado has higher recall (∼0.92) than m6Anet (∼0.51) for m6A detection. Among the overlapping methylated sites between ground truth and computational predictions, there are high correlations of site-specific m6A modification stoichiometry, with correlation coefficient of ∼0.89 for Dorado-truth comparison and ∼0.72 for m6Anet-truth comparison. However, combined assessment of WT and IVT datasets show that while the per-site false positive rate (FPR) can be lower (∼8% for Dorado and ∼33% for m6Anet), both computational tools can have high per-site false discovery rate (FDR) of m6A (∼40% for Dorado and ∼80% for m6Anet) due to the low prevalence of m6A in transcriptome, with a similar trend observed for pseudouridine (∼95% FDR for Dorado). Additional motif analysis reveals that both Dorado and m6Anet exhibit high heterogeneity of false positive calls across sequence contexts, suggesting that sequence contexts help determine accuracy of specific modification calls. There is also a substantial overlap of false positive calls between the two IVT samples, suggesting a post-filtering strategy to improve modification calling by compiling a set of low-confidence sites with a probabilistic model from several IVT samples across diverse cells/tissues. Our analysis highlights key strengths and limitations of the current generation of m6A detection algorithms and offers insights into optimizing thresholds and interpretability. The IVT datasets generated by the RNA004 chemistry provides a publicly available benchmark resource for further development and refinement of computational methods.

## Introduction

RNA modifications are chemical alterations to RNA molecules that extend their functional diversity without altering the underlying genetic code. Most types of RNA carry chemical modifications at some point in their life cycle, which affect stability, structure and RNA–protein interactions[1]. Over 170 distinct types of RNA modifications have been identified across all classes of RNA, profoundly impacting RNA metabolism and function[2]. These modifications were observed to play significant roles in every link of mRNA fate, including pre-mRNA splicing, nuclear export, translation, stabilization and degradation[3]. Recent studies have revealed the important roles of RNA modifying enzymes in various human diseases, including cancer, neurological disorders, cardiovascular diseases, metabolic diseases, as well as developmental and genetic disorders[2, 4]. Revealing the dynamics of these modifications would provide a wealth of information, including modifications and splice sites that are pathogenic, and their consequences at the cellular and organismal levels[5].

Among the diverse array of RNA modifications, nucleobase modifications like N6-methyladenosine (m6A), pseudouridine (Ψ), and 5-methylcytosine (m5C) are the most extensively studied. m6A is the most abundant internal modification in eukaryotic messenger RNA (mRNA) and plays a pivotal role in mRNA splicing, export, stability, and translation efficiency[6]. It is dynamically regulated by “writers” (methyltransferases), “erasers” (demethylases), and “readers” (m6A-binding proteins), and alterations in m6A levels have been implicated in cancer progression, stem cell differentiation, and immune responses[7]. As supported by multiple transcriptome-wide m6A mapping approaches[6, 8], the consensus motif recognized by the m6A writer proteins is best reflected by DRACH, a sequence motif commonly associated with m6A modifications in eukaryotic mRNA[9], where D represents A/G/U, R represents A/G, and H represents A/C/U. Pseudouridine, the isomerized form of uridine, contributes to RNA stability and the accuracy of protein synthesis, and has been associated with ribosomal function and stress responses[10]. m5C is found in various RNA species and is involved in regulating RNA localization, stability, and translation, with links to cancer and other diseases[11].

Detecting and mapping these RNA modifications is essential for understanding their biological functions. Traditional methods for detecting RNA modifications include antibody-based approaches like m6A-seq and MeRIP-seq, which rely on immunoprecipitation of modified RNA fragments[12]. While these methods have provided valuable insights, they suffer from limitations such as low resolution, antibody specificity issues, and inability to quantify modification stoichiometry[13]. Chemical-based methods like PA-m6A-seq and miCLIP offer improved resolution but involve complex procedures and can introduce biases[9]. Enzymatic approaches and mass spectrometry provide high specificity but are often low-throughput or require large amounts of input RNA and sometimes only identify a fraction of the targets[14].

The advent of direct RNA sequencing using nanopore technology has opened new possibilites for detecting RNA modifications at single-molecule resolution. Oxford Nanopore Technologies (ONT) platforms allow RNA molecules to be sequenced directly without reverse transcription or amplification, preserving the native modifications[15]. The computational tools developed to detect RNA modifications from nanopore sequencing data fall under three categories: 1) assessment of basecalling error, 2) statistical comparison of raw signals from two samples, 3) machine-learning-based model inference. Tools like EpiNano[16], Nanom6A[17] and

MINES[18], analyze alignment error profiles in the sequencing reads to infer modification sites. However, these methods do not fully utilize available signal data and can become obsolete due to the ever-developing sequencing and basecalling techniques that continuously reduce error rates. Others, such as Tombo[19], Nanocompore[20], xPore[21] and NanoMod[22], use statistical testing or Bayesian inference to detect changes in the distributions of raw electrical signals between two samples, enabling detection of modification differences between different experimental conditions. This model-free approach can allow *de novo* detection of any type of modification using paired samples, without the need to train a specific model for the modification of interest. However, these methods require control samples to compare modification against, and they do not report modification at single-molecule resolution or per-site stoichiometries.

To overcome these limitations, several deep-learning model-based modification detection tools have been developed. These tools provide models that are trained to detect a specific modification in a query sample without needing a paired control sample. For instance, m6Anet[23] employs a deep neural network trained using multi-instance learning framework to analyze raw current signals and predict m6A modifications with high resolution. RedNano[24] combines raw signal data and error profiles using a deep learning residual network to improve detection accuracy. CHEUI[25] utilizes a two-stage neural network trained on synthetic IVT datasets to enhance reliability. mAFiA[26] advances this by training its model on synthesized short RNA oligonucleotides that replicate sections of actual mRNA containing m6A sites, with precise control over the modification status of each nucleotide.

Despite the increasing number of computational tools for detecting m6A modifications in nanopore dRNA-seq data, their relative performance varies significantly across datasets. A recent benchmarking study systematically evaluated 14 tools for m6A detection using diverse datasets, including synthetic RNA oligos, yeast, mouse, and human transcriptomes[27]. Their results highlighted substantial differences in precision and recall across tools, with multi-condition methods (e.g., Nanocompore, ELIGOS) performing better in high-depth datasets like yeast, while deep-learning-based single-condition tools trained on human transcriptomes (e.g., m6Anet, DENA) excelled in complex transcriptomes like human and mouse. The study also identified biases in certain tools, such as decreased sensitivity in unstructured RNA regions and a tendency for false positives in low-complexity sequences. These findings underscore the importance of careful tool selection and rigorous validation protocols that account for sample complexity, sequencing depth, and RNA structural context.

However, many of these tools were developed using an earlier generation of Oxford Nanopore flow cell chemistry, SQK-RNA002, which is now discontinued. In comparison, the newer flow cell chemistry SQK-RNA004 uses a different motor protein with faster translocation speeds, resulting in a different signal characteristic and higher basecalling accuracy than previous chemistry. As a result, the models trained for modification detection using SQK-RNA002 flow cell cannot be applied to datasets sequenced using the new SQK-RNA004 flow cell. This discrepancy necessitates updating and retraining models to align with the current technology standards. For example, m6Anet has released an updated version (2.1.0) retrained on data from the newest chemistry to improve its accuracy and applicability. Recently, ONT has developed its own base calling and modification detection tool Dorado[28] for SQK-RNA004 flow cell datasets, and claims to be able to detect m5C, m6A, inosine, and pseudouridine (pseU).

In this study, we aim to systematically perform evaluation and calibration of m6Anet[23] and Dorado[28], using in-vitro transcribed (IVT) RNA as a negative control and chemical mapping methods like GLORI[29] and eTAM-seq[30] as ground truth (Figure 1a). While several studies have compared computational tools for m6A detection from ONT direct RNA sequencing, they evaluated obsolete flowcell/chemistry designed for DNA sequencing and focused on approaches that rely on basecalled reads rather than ionic current signals[27, 31]. A recent preprint examined the RNA004 chemistry with Dorado-based detection, and concluded that RNA004 chemistry “significantly improved the throughput, accuracy, and site-specific detection of modifications”[32]. The tools evaluated in our study examine signal information from direct RNA sequencing data for calling m6A or pseU, using two well characterized reference cell lines as well as paired true negative controls. We assessed the ability of these tools to 1) detect transcriptomic sites with m6A methylation, 2) predict m6A stoichiometries that are correlated with ground truth stoichiometries. We use IVT RNA to further examine false positive predictions of Dorado models for pseU detection. Our work provides an analysis of the strengths and limitations of existing tools and seeks to inform future developments in RNA modification detection from direct RNA sequencing data, ultimately contributing to a deeper understanding of the epitranscriptome and its implications for human health.

**Figure 1.**
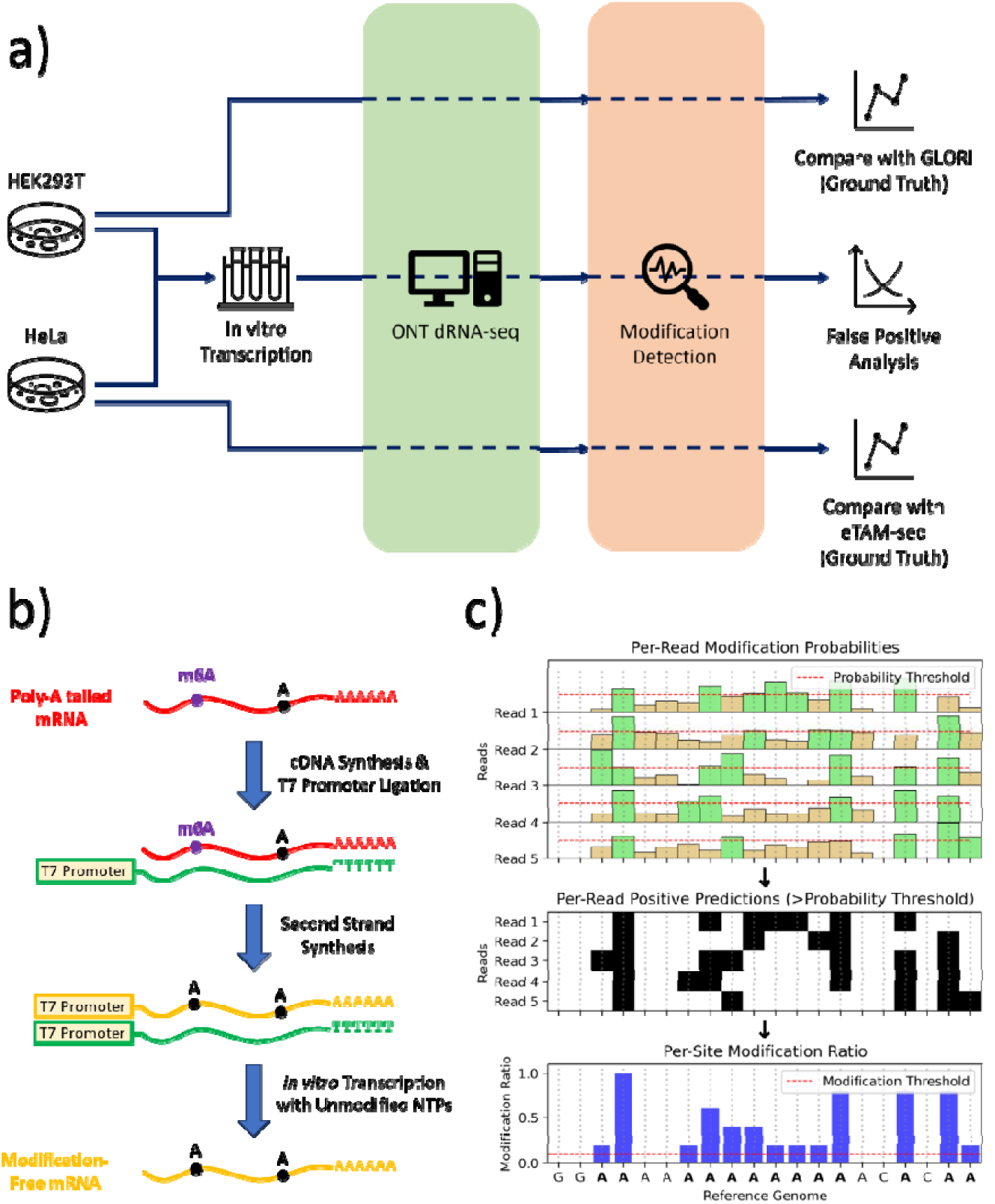
Schematic overview of sample preparation, ONT direct RNA sequencing, and modification detection workflow. a) Study design. Multiple RNA inputs (cell-extracted poly(A)-tailed mRNA containing m6A, or in vitro transcribed controls without modifications) are processed through Oxford Nanopore’s direct RNA sequencing platform. The resulting signal and basecalls then feed into a modification detection module, which assigns per-read modification probabilities. b) The preparation of cellular m6A-modified mRNA (red), in vitro transcripts (green), and reverse-transcribed unmodified transcripts (yellow). c) Schematic demonstrating how per-read probabilities are filtered by a chosen threshold to produce positive modification calls (green and black), which can then be aggregated into per-site modification ratios across the transcriptome, and further judged by certain threshold (10% in this study) for site-level predictions.

## Results

### Summary of the experimental design

For evaluating m6A detection, we performed direct-RNA ONT sequencing of mRNA from wild-type (WT) HEK293T and HeLa cell lines, and compared them against m6A ground truth available from GLORI[29] and eTAM-seq[30], respectively. For further evaluation of false positive predictions, we performed in-vitro transcription (IVT) of mRNA from wild-type HEK293T and HeLa cell lines using standard NTPs to obtain modification-free RNA (**Figure 1**b). Both the WT mRNA and the IVT RNA were sequenced using the Oxford Nanopore Direct RNA SQK-RNA004 kit with separate PromethION RNA flow cells (**Figure 1**a). The overall expression of transcript isoforms in the IVT RNA and wild-type mRNA were strongly correlated in both and cell lines (Supplementary Figure 1). General information and quality control for the raw sequencing data are provided in Supplementary Table 1.

We will elaborate on the intricate technical details of RNA modification here. Dorado and m6Anet predict the probability of m6A modification for each candidate adenosine in a read (that is, per-read modification probability) and predict it to be modified if the probability is above a certain threshold. The default probability threshold for m6Anet is 0.033, whereas Dorado relies on Modkit to determine the optimal threshold by sampling the distribution of probability predictions (generally ∼0.61 from our experience, see Supplementary Table 2 for details).

Dorado assesses all adenosines in a read for m6A modification and the reads can be aligned to a reference genome using the splice-aware alignment mode of minimap2[33]. Then, the “pileup” module in Modkit calculates the modification ratio for all DRACH sites in the reference genome (that is, per-site modification ratio), which is calculated as the number of m6A predictions mapped to the reference DRACH site divided by the total number of A’s mapped (modified or unmodified). However, Modkit uses separate thresholds for modified and unmodified predictions by discarding intermediate or low confidence probability calls. This complicates the performance assessment of a binary classification model at different probability decision thresholds. To address this challenge in our comparative evaluation study, we calculated the per-site ratio for Dorado by tabulating per-read output from Modkit’s “extract” module instead of the “pileup” module, without discarding intermediate probability calls.

On the other hand, m6Anet aligns a read sequence (via minimap2) and the corresponding raw signal (via f5C) to spliced reference transcript sequences. Then, it assesses whether the chunk of signal aligned to a reference transcriptome DRACH site represents an unmodified A or m6A nucleotide. m6Anet also requires a minimum read coverage (20 by default) at a DRACH site to assess the reads at that site for methylation. The ground truth for m6A provides modification ratios for genomic sites only, but a DRACH site in the reference genome may correspond to DRACH sites in multiple transcripts of a gene. Therefore, we mapped the transcript coordinates of m6Anet predictions back to genomic coordinates for evaluation, as shown in Supplementary Figure 2. In the following sections, “per-read modification probability” prediction refers to single-molecule and single-base probability of modification predicted by deep-learning models of Dorado and m6Anet. On the other hand, “per-site modification ratio” prediction refers to the fraction per-read predictions classified as modified for a given genomic site (**Figure 1**c).

Furthermore, we used several performance measures in our evaluation study (see Methods for details). In addition to commonly used measures such as true positive (TP), false positive (TP), false negative (TN), we also calculate the false positive rate (FPR) in a per-read and per-site basis. Following the strategies used for evaluating DNA modification prediction, we also assessed a false discovery rate (FDR) measure [34]: unlike its canonical meaning in hypothesis testing, in the context of this study, FDR is calculated as the ratio of the fractions of modified predictions in IVT and WT samples.

### Evaluating Per-Site Recall and Molecular Stoichiometry of Predicted m6A Sites

For this evaluation, a reference DRACH site is considered a positive per-site prediction for a tool if the modification ratio or stoichiometry is ≥10% ; as mentioned earlier, modification ratio is defined as “the number of m6A per-read predictions with modification probability above a fixed threshold” divided by “the total number of A’s” at the site (**Figure 1**c). We chose 10% as the modification ratio cut-off for classifying as site as methylated because the ground truths for HEK293T and HeLA used this cut-off value. A positive per-site prediction in wild-type samples is considered a true positive prediction if it is present in the ground truth; the evaluation strategy is described in further detail in the Methods section. Since the ground truths for HEK293T (GLORI) and HeLa (eTAM-seq) report only the sites that have m6A modification ratio ≥10%, it is not possible to reliably determine false positive calls for wild-type samples, thus prompting the need for IVT RNA. We restricted GLORI and eTAM-seq ground truth to DRACH sites that have ONT sequencing coverage ≥10 to prevent coverage differences between sequencing platforms from effecting the evaluation of deep-learning models.

For both algorithms, WT samples consistently show higher modification rates than the corresponding IVT controls, reflecting the expected presence of true modifications. Nevertheless, the IVT samples retain a background signal—indicating false positive calls, which we explore further below. We also observed a positional bias along the transcript, underscoring that modification rates vary systematically with read position. Notably, both Dorado and m6Anet show elevated calls near the 3′ end of the reads (Supplementary Figure 3).

For HEK293T cell line, Dorado and m6Anet produced predictions for ∼0.5M and ∼0.3M sites, respectively, out of which 12.8% and 13.2% overlap with the ground truth. Almost all sites analyzed by m6Anet are covered by Dorado, and almost all sites reported in the ground truth datasets are covered by Dorado, as shown in Figure 2a. Similar observations are made for the prediction on the HeLa cell line. We note that since m6Anet relies on f5c to perform signal alignment to spliced transcript sequences, it only considers positions that fall within annotated transcript regions with sufficient read coverage, restricting its ability to detect m6A sites located outside or beyond well-characterized transcripts, as described in Supplementary Figure 2. This ultimately reduces the number of discoverable sites compared to methods that utilize genome-based alignments. Thus, incorporating read alignments to transcript sequences within modification calling can allow the uncertainties and errors of read-to-transcript assignment to negatively affect modification calling.

**Figure 2:**
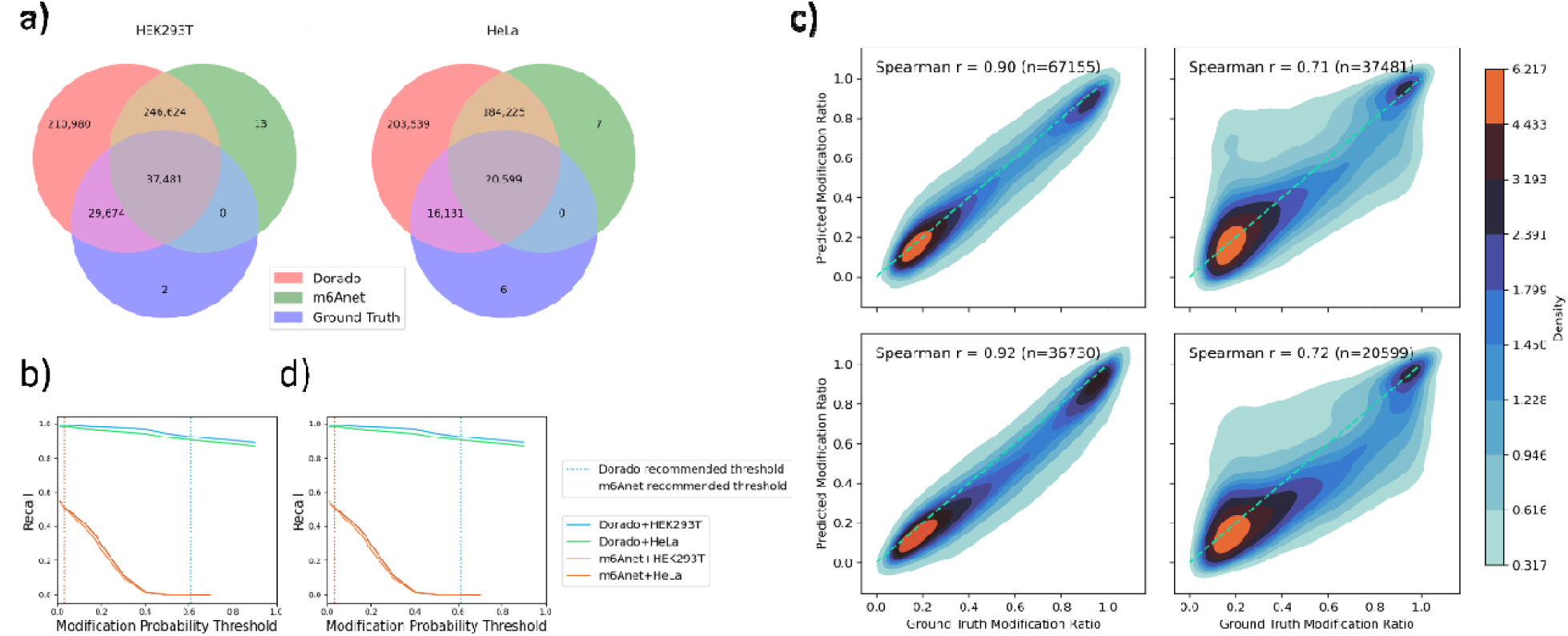
m6A detection performance comparisons between Dorado and m6Anet in HEK293T and HeLa datasets. a) Venn diagrams depicting the overlap among Dorado predictions, m6Anet predictions, and ground truth sites for HEK293T (left) and HeLa (right). Each circle segment denotes the number of exonic DRACH sites predicted by the respective method or ground truth. Note that these sets are the counts of the raw outputs before called for modification. b,d) Per-site recall (left) and modification ratio correlation (right) curves plotted against the modification probability threshold for Dorado (green and blue lines) and m6Anet (red and orange lines) on both HEK293T and HeLa datasets. Vertical dotted lines indicate each model’s recommended threshold, illustrating how recall and correlation vary as the threshold changes, and highlighting the trade-offs each tool makes between sensitivity (recall) and predictive reliability (correlation). c) Density plots illustrating correlations between predicted modification ratios from Dorado (left column) or m6Anet (right column) and the ground truth modification ratios for HEK293T (top row) and HeLa (bottom row). The Spearman correlation coefficients are shown in each panel, along with the total number of sites (n). Dense regions (red) indicate a strong agreement between predicted and ground truth ratios, while lighter areas (blue) reflect lower site densities or greater deviations.

To evaluate Dorado and m6Anet’s ability to detect sites known to have m6A methylation, we calculated per-site recall at various per-read probability thresholds ranging from 0 to 1, including the default probability thresholds (Figure 2b). As the probability threshold increases, recall decreases across all conditions as expected. However, Dorado and m6Anet diverge significantly in their optimal threshold regions. Dorado maintains relatively high recall across both HEK293T and HeLa datasets, indicating a stable performance that does not drop sharply when adjusting the probability threshold. By contrast, m6Anet starts with decent recall but quickly loses sensitivity as the threshold is raised, which suggests a more threshold-sensitive approach. These observations align with each tool’s recommended thresholds (Dorado: ∼0.61; m6Anet: ∼0.033). However, even at the recommended modification probability threshold for m6Anet, the recall is 51.5%, with 90% false negative rate arising from no predictions made and 10% false negative rate arising from incorrect predictions, while for Dorado the recall is 92.5% with almost no false negative rate arising due to no predictions made (<0.01%). The default threshold numbers for Dorado are estimated with slight variations by ModKit depending on the sample, and the actual numbers are shown in Supplementary Table 2. At these thresholds, recall remains relatively high. Thus, the results underscore that each model’s recommended threshold provides a distinct balance of sensitivity, and adhering to these model-specific thresholds generally leads to optimal or near-optimal performance. Additional metrics and numbers at suggested threshold are presented in Supplementary Table 3.

Evaluation of recall only assesses the ability to predict presence or absence of m6A methylation at a site as a binary outcome using ≥10% modification ratio. However, in real-world applications, examining differential methylation between two experimental conditions or samples requires measuring statistically significant changes in methylation stoichiometries, as opposed to determining the mere presence or absence of methylation. Therefore, it is important to evaluate the accuracy of methylation stoichiometries predicted by each method. For this evaluation, we calculated the Spearman correlation between the predicted and ground truth methylation ratios at the true positive sites of each tool using default or recommended per-read modification probability thresholds. **Figure 2**c shows the heatmap plot of predicted and ground truth per-site modification ratios. Under the recommended per-read modification probability threshold, Dorado achieved a Spearman correlation coefficient around 0.9 for both cell lines, while m6Anet had coefficients just over 0.7. The strong correlations suggest that both models have captured the extent of m6A modifications faithfully, while Dorado also demonstrates its excellent performance in predicting actual stoichiometry values. The heatmap also suggests that both Dorado and m6Anet have a slight tendency to underpredict the modification ratios relative to ground truth (**Figure 2**c, according to the diagonal reference line). Figure 2d illustrates how the correlation with ground truth changes as the per-read modification probability threshold shifts, generally mirroring the trend seen in the recall curve. However, one notable deviation is a substantial peak early in m6Anet’s correlation—around a threshold of 0.1—which does not align with its recommended threshold of 0.033. This indicates that while m6Anet can achieve high correlation at low per-read modification probability thresholds, that optimal threshold setting in this case differs slightly from the one officially recommended for both cell line samples. Meanwhile, a side-by-side comparison of Dorado’s and m6Anet’s predicted per-site modification ratios (Supplementary Figure 4) reveals that both overpredicted and underpredicted sites contribute to the differences between the two tools, with m6Anet tending to overpredict in general.

### Evaluation of False Positive m6A Predictions

The available ground truth for wild-type (WT) HEK293T and HeLa cell lines only includes sites with a high confidence on the presence of m6A methylation and does not include sites with a high confidence on the absence of m6A methylation. Therefore, it is not possible to reliably determine false positive calls and examine the precision of these modification detection models using WT samples. To address this, we performed in-vitro transcription (IVT) of HEK293T and HeLa WT cell lines using canonical NTPs to obtain modification-free RNA. This makes IVT RNA an ideal resource for evaluating false positive predictions of Dorado and m6Anet at both per-read and per-site resolution, since any m6A prediction in IVT RNA is a false positive. We assessed the false positive predictions from IVT samples using per-read and per-site false positive rate and false discovery rate.

Per-read false positive rate (per-read FPR) for IVT RNA is defined as the total number of m6A predictions above a certain per-read modification probability threshold (i.e., false positive predictions) divided by the total number of predictions in the sample. Similarly, per-site false positive rate (per-site FPR) is defined as the number of sites with ≥10% modification ratio divided by the total number of sites. **Figure 3**a shows how the per-read false positive rate of IVT samples changes as the per-read modification probability threshold changes, as well as the corresponding per-site recall and correlation of WT samples at those thresholds as a reference. The results for HEK293T and HeLa (**Figure 3**a, left and right) are very consistent. At their respective recommended thresholds, m6Anet—although demonstrating a more stringent false positive rate (FPR) over much of the threshold spectrum—must lower its cutoff dramatically to have reasonable recall and correlation, ultimately pushing up its per-read FPR (∼10% at recommended threshold), higher than Dorado’s. In contrast, Dorado retains consistently strong recall and correlation across various modification probability thresholds, allowing it to use a higher cutoff; as a result, its per-read FPR at the recommended threshold (∼0.61) is 3%, lower than m6Anet’s.

**Figure 3:**
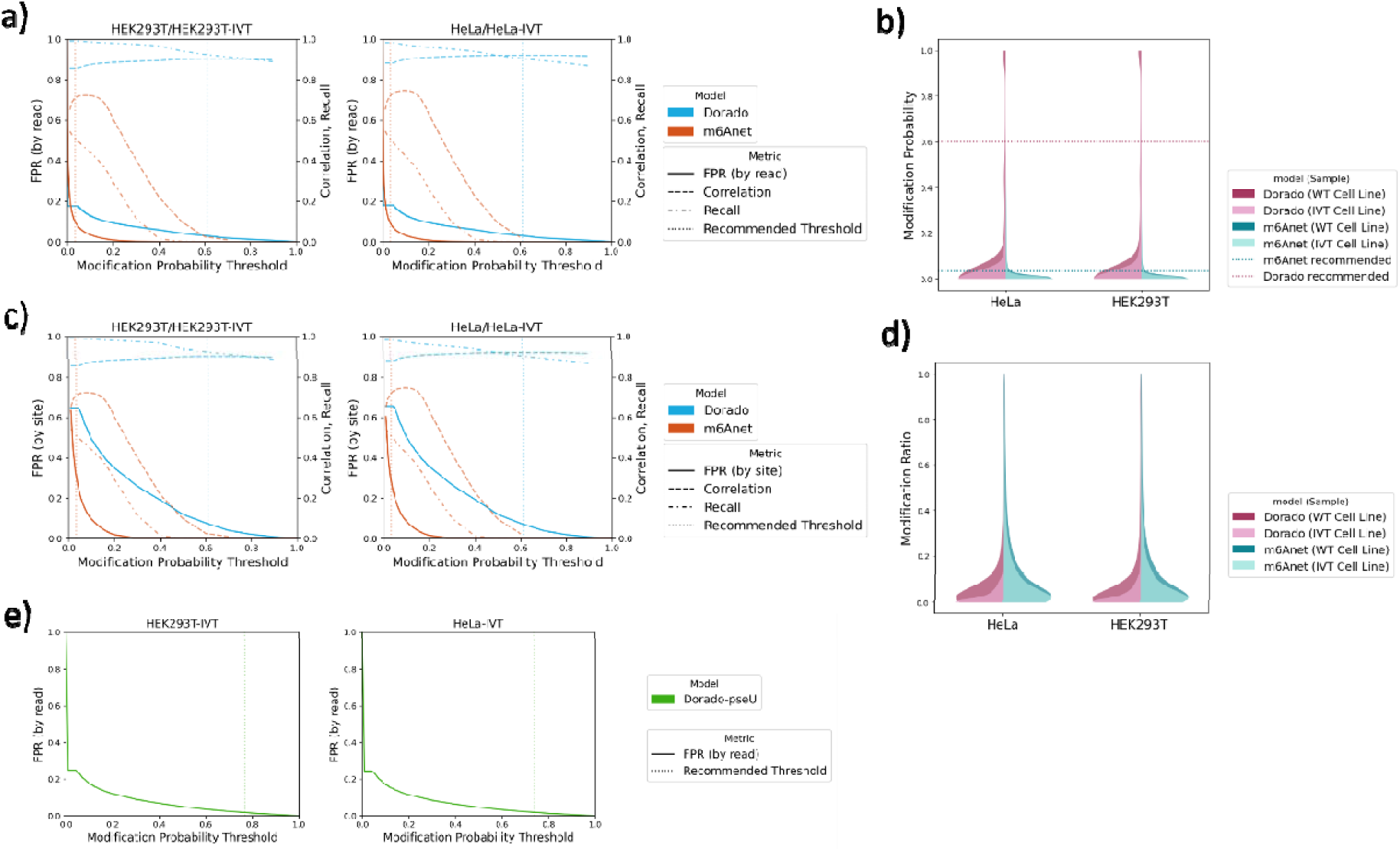
Threshold-dependent performance and distribution of predicted m6A probabilities/ratios for Dorado and m6Anet in HeLa and HEK293T datasets. a) Per-read false positive rate (FPR), with per-site recall, and correlation as functions of the modification probability threshold in HEK293T (left) and HeLa (right) (using IVT control for FPR, WT RNA for correlation and recall). The dotted vertical lines indicate Dorado’s (blue) and m6Anet’s (orange) recommended thresholds. c) Per-site FPR, recall, and correlation under the same settings as a). b) Violin plot showing the distribution of per-read modification probabilities for wild-type and IVT samples (HeLa and HEK293T), with horizontal dotted lines marking each model’s recommended threshold. Notably, Dorado’s predictions display a more pronounced double-peaked structure split at its recommended threshold, while m6Anet’s distributions exhibit a single peak closer to zero. d) Violin plots present the corresponding per-site modification ratio distributions under the same settings as c). e) Per-read false positive rate (FPR) of Dorado’s pseudouridine model, as a function of the modification probability in two cell lines. The dotted vertical lines indicate Dorado’s recommended thresholds.

We examined the distribution of per-read modification probabilities from a different perspective in Figure 3b. Dorado’s predictions exhibit a distinct double-peaked pattern, with its recommended threshold neatly dividing the low- and high-probability peaks at each end of the range. By contrast, m6Anet’s predicted probabilities do not show a comparable bimodal structure, and its recommended threshold primarily functions as a cutoff for lower-probability predictions. This contrast in distributions reflects each tool’s distinctive predictive characteristics and supports the notion that Dorado’s recommended threshold more clearly partitions truly modified bases from unlikely ones.

The per site evaluation of the FPR (**Figure 3**c) shows a similar result as the per read FPR, and comparing the distribution of modification ratios between WT and IVT samples (**Figure 3**d) also supports that Dorado is better able to distinguish the modified or unmodified sites. At Dorado’s recommended threshold (∼0.61), the per-site FPR for HEK293T is ∼7.7%. At m6Anet’s recommended threshold (∼0.033), the per-site FPR rises to ∼33.3%. Thus, while m6Anet can capture more true modifications at lower thresholds compared to higher thresholds, it also inflates the number of false positives. Dorado’s threshold, by contrast, yields fewer false positives, underscoring each model’s distinct balance between sensitivity and specificity at their recommended cutoffs.

### Using False Discovery Rate to Estimate the Reliability

Evaluation of the false positive rate tells us the percentage of unmodified A nucleotides or sites that will be incorrectly predicted as modified. However, if the true modification has low abundance in a sample (that is, for rare modifications), then even a small false positive rate can overwhelm the overall positive predictions. A modification can have low abundance due to low modification ratio on average or due to low number of sites where the modification is found irrespective of the modification ratio. There are more than 8M exonic DRACH sites in the human reference genome GRCh38, but only 170k (1.9%) and 70k (0.8%) of these sites were detected to have m6A methylation with at least a 10% modification ratio in the ground truth for HEK293T and HeLa, respectively. Moreover, even among the sites with at least a 10% modification ratio, the median modification ratio is 39% and 48% in HEK293T and HeLa, based on the ground truth. Therefore, even a 5% per-site FPR from Dorado or m6Anet could result in false positive predictions dominating overall positive predictions. To further assess this, we followed an evaluation strategy devised by Kong et al [34] for estimating the false discovery rate (FDR) using modification free sample, as described in Methods. Briefly, we estimate per-read false discovery rate as the fraction of per-read predictions in IVT sample predicted as modified (i.e. modification probability above a certain threshold) divided by the fraction of predictions in WT samples predicted as modified. Similarly, per-site false discovery rate is the fraction of sites in IVT sample predicted as modified (i.e., modification ratio ≥10%) divided by the fraction of sites in WT samples predicted as modified. The idea behind this strategy is that positive predictions of WT samples can be decomposed into true positives and false positives, where the latter can arise 1) due to the confounding effect of nearby modifications or 2) due to inherent signal noise or model biases (e.g. overfitting) even when no modification is present. Positive predictions in IVT can be used to estimate the abundance of the second type of false positive predictions that can arise in a WT sample. Thus, the proposed FDR provides a lower bound estimate of the true FDR of wild-type samples.

Figure 4a and b illustrate how the per-read and per-site false discovery rate (FDR) in IVT samples (HEK293T and HeLa) changes across a range of per-read probability thresholds, alongside the corresponding per-site recall and correlation in the WT samples. In the per read evaluation, Dorado’s per-read FDR remains consistently above m6Anet’s, while m6Anet plateaus at a certain FDR (∼0.25) level in both cell lines, implying that its rate of false calls does not improve further beyond a particular modification probability threshold. By contrast, the persite evaluation shows that for HEK293T, m6Anet’s per-site FDR is over 84% at default modification probability thresholds but can drop substantially at high thresholds, whereas Dorado’s FDR is over 32% at default thresholds and remain high at higher thresholds.

**Figure 4:**
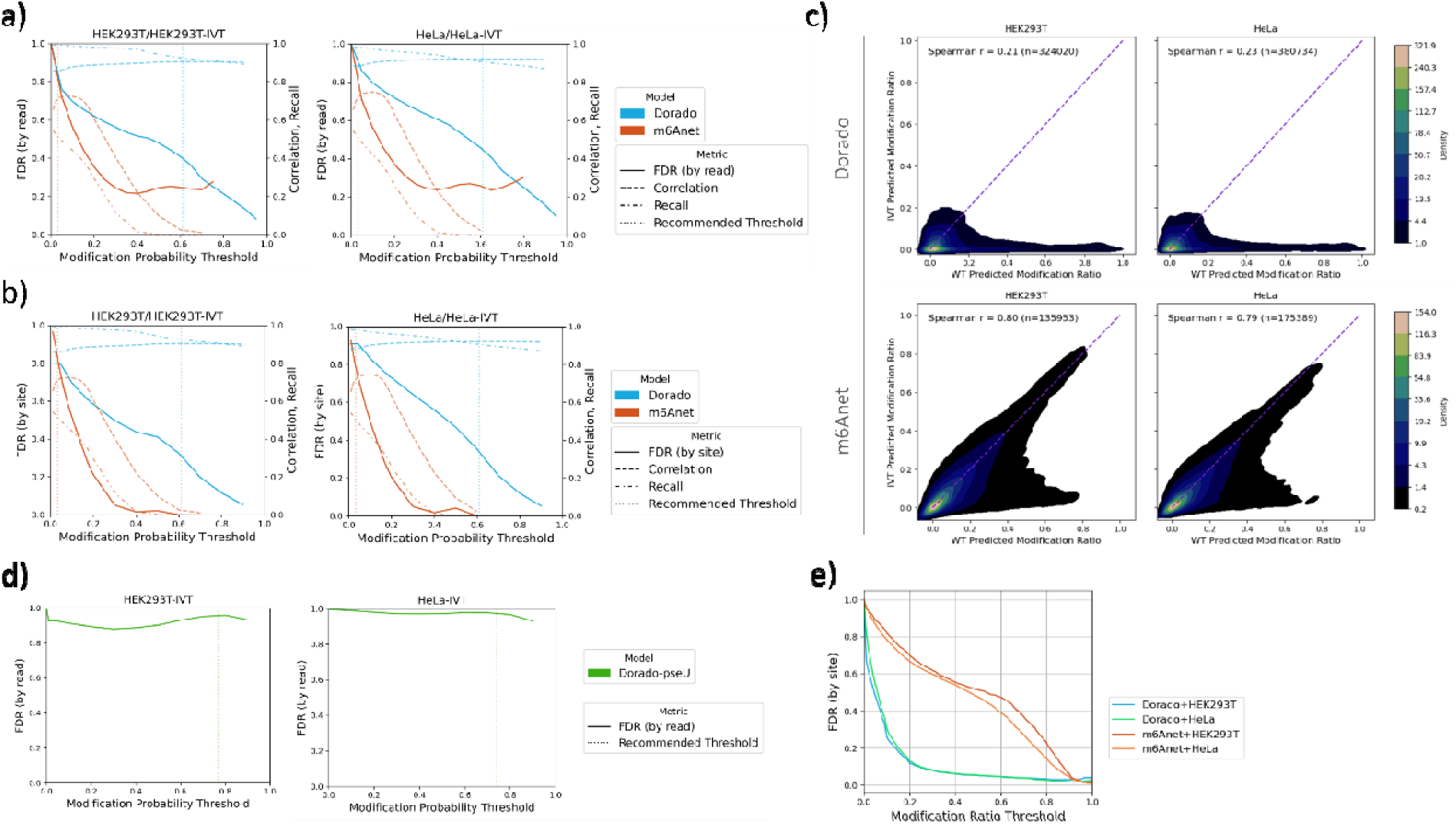
Threshold-dependent false discovery rate (FDR), recall, and correlation analyses for Dorado and m6Anet in HEK293T and HeLa datasets. a) shows how the per-read FDR (solid lines), and per-site recall (dash-dotted lines) and correlation (dashed lines) change as the modification probability threshold varies for each model. b) presents analogous curves for the per-site FDR. The vertical dotted lines indicate each model’s recommended threshold. FDR calculation at very high threshold is removed to avoid noise effect due to the low absolute number of positive predictions after filtering (>10). c) Comparison of per-site modification ratios in IVT (y-axis) versus WT (x-axis) samples for Dorado (top row) and m6Anet (bottom row) in HEK293T (left column) and HeLa (right column). The Spearman correlation coefficients (r) in each plot quantify the degree of similarity between the IVT and WT predicted modification ratios. Dorado’s correlations (0.18–0.24) indicate limited agreement. By contrast, m6Anet shows substantially stronger IVT–WT correlations (0.79–0.80). d) Per-read false discovery rate (FDR) of Dorado’s pseudouridine model, as a function of the modification probability in two cell lines. The dotted vertical lines indicate Dorado’s recommended thresholds. e) Per□site FDR as a function of the modification ratio threshold for Dorado and m6Anet in HEK293T and HeLa datasets. As the threshold increases, sites must exhibit increasingly higher modification ratios before being labeled “modified,” which generally reduces the FDR.

We also compared Dorado and m6Anet’s predicted per-site modification ratios in IVT versus WT samples for both cell lines (Figure 4c). Dorado’s result reveals low overall correlation (Spearman’s r = 0.21–0.23). Most sites cluster near (0,0), as expected for unmodified bases, yet we observe a sparse portion of points at or near zero in the WT sample but non-zero in the IVT sample—indicating false positives sites in the IVT dataset. This pattern underscores that some sites flagged by Dorado as moderately to be modified in IVT are predicted to have negligible modification in WT. Moreover, the lower left region of the plot reflects a non-trivial fraction of shared sites that show intermediate modification ratios in both samples. In fact, 10% (HEK293T) and 9% (HeLa) of these sites exhibit modification ratios between 5% and 40% in both IVT and WT, raising the possibility of model biases or systematic errors that yield moderate modification signals in both modified and unmodified RNA. By contrast, m6Anet shows a substantially higher IVT–WT correlation (Spearman’s r = 0.79–0.80), as illustrated in the lower panels of Figure 4c. 32% (HEK293T) and 30% (HeLa) of these sites exhibit modification ratios between 5% and 40% in both IVT and WT. This tighter clustering along the diagonal suggests that m6Anet assigns more similar modification ratios to corresponding sites in IVT and WT, potentially reflecting that the model is systematically labeling many sites with moderate to high modification probabilities regardless of whether the sample has modifications or not. Thus, m6Anet may be overfitted to some extent to capture background noise or intrinsic biases that produce artificially elevated modification ratios.

Another question we next address based on the FDR is of how reliable high modification ratio calls are, and how high a ratio must be before a site can be considered confidently modified. Fixing the per-read probability threshold at recommended values, we observe that both FDR for Dorado and m6Anet starts near 100% and then declines as the threshold increases, as shown in Figure 4e. Notably, Dorado’s curves drop more sharply at moderate thresholds—indicating that for sites with modestly high modification ratio (>20%), the likelihood of it being a false call decreases substantially. By contrast, m6Anet maintains higher FDR at moderate thresholds but eventually converges on similarly low FDR at stricter cutoffs.

From a practical perspective, these trends imply that sites with higher modification ratios are generally more trustworthy. However, the exact threshold at which FDR becomes acceptable depends on the specific model, as well as the user’s tolerance for false positives. For instance, researchers using Dorado aiming for high specificity could require a site to surpass a 0.2 modification ratio, accepting fewer overall calls but more reliable ones, while for m6Anet they would instead only focus on the sites with modification ratio greater than 0.8. Conversely, lowering the threshold can capture more potentially modified sites at the risk of inflating FDR. Ultimately, tuning the per-site modification ratio threshold based on desired FDR can balance sensitivity and specificity, helping users decide how to classify “high-confidence” modified sites in their datasets.

Figure 3 In summary, our evaluation of false positive m6A predictions highlights distinct performance characteristics of Dorado and m6Anet. Each model demonstrates unique strengths and trade-offs in balancing sensitivity and specificity, influenced by their predictive thresholds and internal architectures. While Dorado benefits from a more distinct partitioning of true modifications from noise, m6Anet exhibits improved specificity but faces challenges in maintaining sensitivity at stringent thresholds to ensure its prediction power. False discovery analyses further reveal systematic biases and context-dependent behaviors in both models, underscoring the complexities of RNA modification detection. These findings provide a nuanced understanding of the models’ predictive capabilities and limitations, offering valuable insights for their application in different experimental contexts.

### Evaluation of False Positive Pseudouridine Predictions

Since Dorado also provides models for pseudouridine modification, we also analyzed the false positive rates using the IVT samples (Figure 3e). The overall shape of the pseudouridine FPR curve is similar to what we observed with m6A models: at very low thresholds, the model calls many reads as modified, resulting in a sharp initial drop in FPR as the threshold increases. However, the pseudouridine model achieves near-zero FPR at higher thresholds—close to its recommended settings. Under suggested thresholds, the FPR based on the IVT samples are 1.77% and 1.85% for HEK293T and HeLa, respectively. However, we caution that due to the extremely low expected pseudouridine levels in human transcriptome, as reported previously that only 2209 confident pseudouridine sites were revealed in HEK293T cells[35], the FDR could still be very high. Figure 4d illustrates how this issue manifests in both HeLa and HEK293T data, showing a consistently close to FDR of 100%, underscoring—despite near-zero FPR at tighter thresholds—pseudouridine calls must be approached cautiously. Additional orthogonal validation or complementary sequencing strategies may be needed to confirm high confidence pseudouridine sites in human samples.

### Examining RNA Modification Differences Between HEK293T and HeLa

A natural question arising from our analysis is how modification profiles differ across cell lines. To address this, we compared the per-site modification ratios between HeLa and HEK293T and their corresponding IVT samples, as well as the ground truth datasets, on the common sites predicted by Dorado and m6Anet (Figure 5) with ONT sequencing coverage >10.

**Figure 5:**
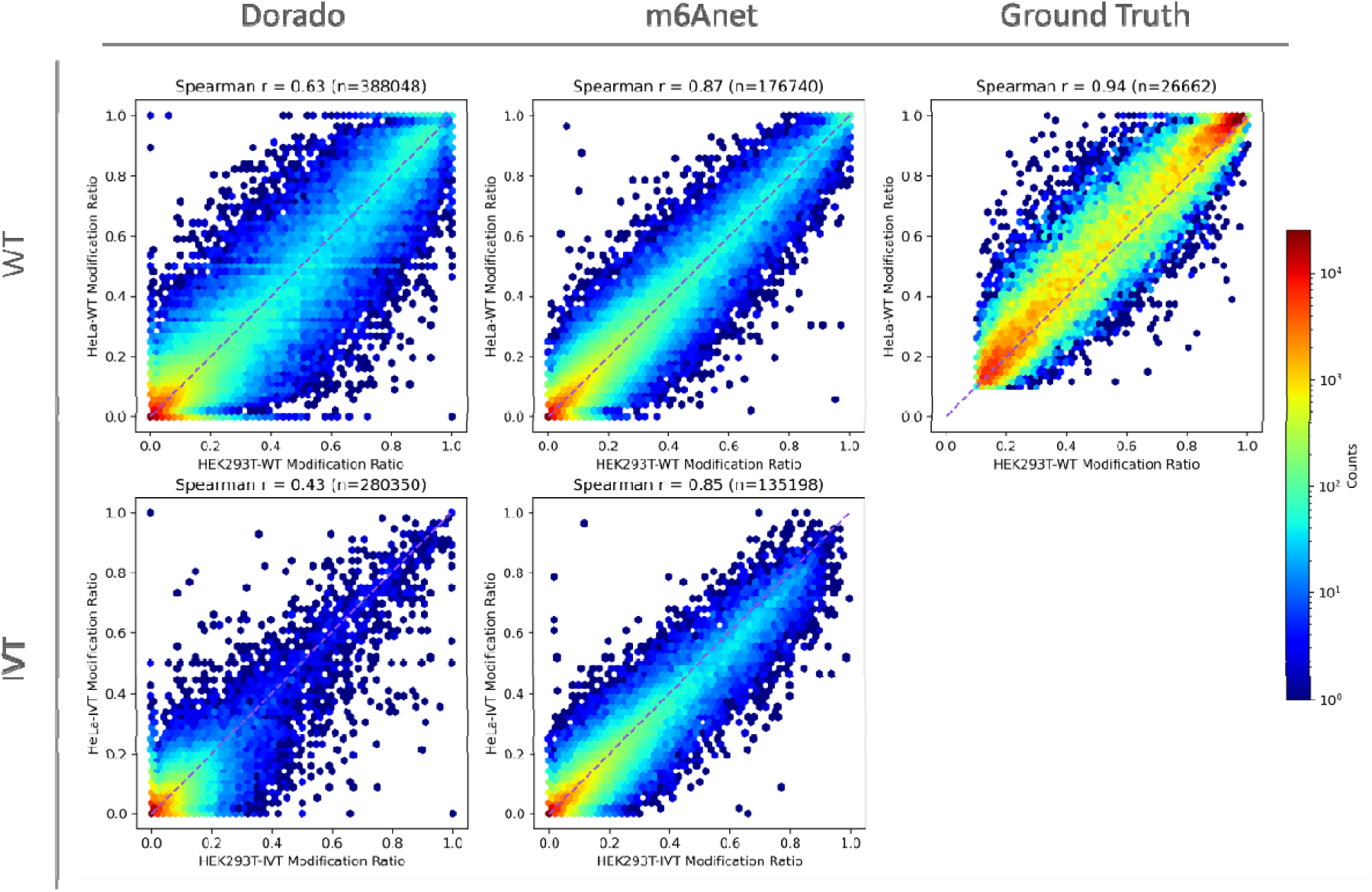
Comparison of modification ratio correlations between HEK293T and HeLa in both WT and IVT samples for Dorado (left), m6Anet (middle), and ground truth (right). In the top row, each hexbin plot shows the per-site modification ratios for HEK293T-WT (x-axis) against HeLa-WT (y-axis); in the bottom row, the ratios for HEK293T-IVT (x-axis) are compared to HeLa-IVT (y-axis). The Spearman correlation coefficients (r) are noted above each panel, along with the number of common sites (n). For Dorado, we observe moderate cross-cell-line correlation in WT samples and lower correlation in IVT. By contrast, m6Anet yields a higher correlation in both WT and IVT, suggesting stronger internal consistency across samples. Ground truth shows the highest correlation. The color scale (right) indicates the log-scale count of sites within each hexbin.

For Dorado, we identified 388,048 common sites in the two WT samples, representing ∼74% of HEK293T calls and ∼91% of HeLa calls, with a Spearman correlation of 0.63 for stoichiometry— indicating a moderate level of agreement in modification levels between the two cell lines. For m6Anet, we found 176,740 common sites, representing ∼62% of HEK293T calls and ∼86% of HeLa calls, with a higher correlation of 0.87, suggesting a stronger cross-cell-line concordance. In contrast, as expected, Dorado shows substantially lower overlap and correlation for the IVT samples of the two cell lines, while m6Anet maintains an unexpectedly strong correlation even in IVT. This discrepancy hints that m6Anet may be capturing an internal, genome-wide pattern rather than directly reflecting truly modified versus unmodified bases, raising the possibility that its predictions are driven more by inherent model biases than by genuine m6A signals. Moreover, the substantial overlap of false-positive calls between the two IVT samples in both models suggests a potential strategy of compiling a “blacklist” of recurrently miscalled sites. If constructed using multiple IVT datasets from diverse cell types (to ensure coverage of diverse transcripts), such a probabilistic blacklist might be able to serve as an effective post-filter to further improve the specificity of RNA modification calls in direct RNA sequencing.

### Examining the Impact of Motif Context on m6A Detection

Understanding the motif context of m6A modifications provides insight into how specific sequence patterns influence a model’s ability to detect true sites while minimizing false positives. A detailed look at motif-specific performance reveals substantial variation in correlation, recall, FPR, and FDR across different 5Dmer contexts in DRACH motifs, underscoring the significant impact of sequence features on m6A detection (Figure 6).

**Figure 6:**
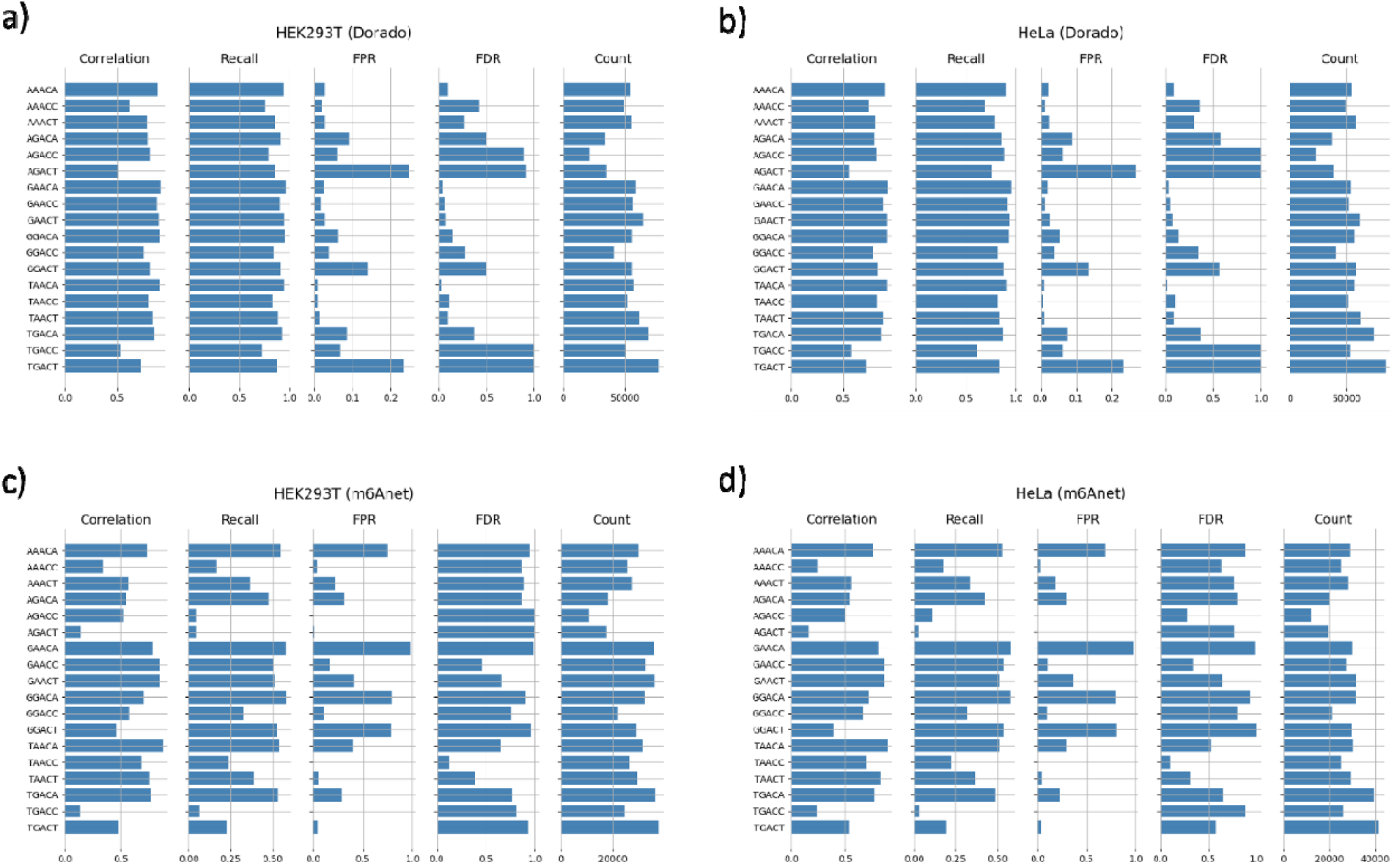
Motif-Specific Performance Metrics for Dorado and m6Anet in HEK293T and HeLa. Each panel displays bar plots of correlation, recall, FPR (per site), FDR (per site), and count (i.e., the number of sites matching each motif) for different 5-mer motifs around the predicted adenines. The top row shows Dorado results in HEK293T (a) and HeLa (b), while the bottom row presents m6Anet results in HEK293T (c) and HeLa (d). As defined earlier, correlation measures how well the predicted modification ratios match the ground truth; recall (sensitivity) is the fraction of truly modified sites correctly identified; FPR is the fraction of unmodified sites (in IVT) misclassified as modified; and FDR is the fraction of predicted positives between IVT and WT (capped at 1). Comparing these metrics across different motifs highlights that sequence contexts greatly influence the accuracy of modification detection.

Comparing HEK293T and HeLa side by side shows that both Dorado and m6Anet exhibit consistent trends across cell lines, suggesting their motif-associated biases are largely independent of cell type. However, Dorado demonstrates a notable repeating pattern in its false positive and false discovery rates when sorting the motifs by alphabetical order (Figure 6a, b), implying a bias wherein specific nucleotides or positions within the motif trigger higher misclassification rates. Additionally, m6Anet exhibits even stronger motif-dependent heterogeneity, with some motifs having virtually no false discovery and others being dominated by false calls, yet similar motifs do not have similar false positive rates when sorted alphabetically (Figure 6c, d). These observations highlight the critical role of sequence context in current m6A detection models and suggest that careful consideration of motif features could guide future refinements to improve algorithmic calibration and performance.

## Methods and Materials

### Benchmark Datasets

The ground truth datasets were obtained from the GLORI[29] and eTAM-seq[30], studies, providing chemically detected, genome-wide m6A modification atlases for HEK293T and HeLa cell lines, respectively. For GLORI, we downloaded the summarized m6A sites identified in HEK293T cells from the supplementary data, which contains the measured m6A levels for each genomic site across two technical replicates. We took the mean value of the two replicates and select only the exonic positions, resulting in (n = 142892) total modified sites. For eTAM-seq, we accessed the transcriptome-wide profiling and quantification of m6A in HeLa cells (GSE201064), which includes methylation levels from three technical replicates and one deeply sequenced replicate in combination with IVT. To ensure consistency, we united the replicates and calculated the mean m6A level for each genomic site (n = 64272). All these recorded sites were further filtered on their corresponding coverage in our ONT sequencing (ONT sequencing coverage>=10) to reduce the noise in the evaluation. These datasets serve as benchmark, chemically validated ground truth references for m6A modifications at base resolution and were used to evaluate the accuracy of RNA modification detection models.

### IVT and Sequencing Datasets

Total RNA from the HeLa cell line was purchased from Takara Bio (Cat. No. 636543), while total RNA from HEK293T cells was extracted by our collaborators at the University of Pittsburgh (UPitt). Before in vitro transcription (IVT), cDNA was generated from both RNA samples to incorporate the T7 RNA polymerase promoter sequence, allowing binding of the RNA polymerase to the cDNA. Three primers were used in this process, all ordered from IDT: a T7 template-switch oligo (T7-TSO; 5’-ACTCTAATACGACTCACTATAGGGAGAGGGCrGrG+G-3’), an oligo-(dT) primer (5’-TTTTTTTTTTTTTTTTTTTTTTTTTTTTTTVN3’), and a T7 extension (T7-ext) primer for PCR (5’-GCTCTAATACGACTCACTATAGG-3’). mRNA was isolated from the total RNA using the NEBNext® High Input Poly(A) mRNA Isolation Module (NEB, E3370S) following the manufacturer’s protocol.

For cDNA synthesis, 200 ng of mRNA was mixed with 7.5 µL of water, 2.5 µL of 2.5 µM oligo-(dT) primer, and 1.0 µL of 10 mM dNTPs. This mixture was incubated at 75°C for 3 minutes, followed by 42°C for 2 minutes, and then snap-cooled on ice. A strand-switching mixture was prepared by adding 4 µL of 5x RT buffer (ThermoFisher, EP0752), 1 µL of RNaseOUT (Invitrogen, 10777019), 2 µL of 10 µM T7-TSO primer, and 1 µL of nuclease-free water to the previous mRNA mixture, which was then incubated at 42°C for 2 minutes. After this incubation, 1 µL of Maxima H Minus Reverse Transcriptase (ThermoFisher, EP0752) was added, and reverse transcription with strand-switching was performed under the following conditions: 42°C for 90 minutes, 85°C for 5 minutes, and held at 4°C. To digest the RNA template, 1 µL of RNase cocktail enzyme mix was added to the reaction. Clean-up was performed using a 0.8x bead ratio, followed by two washes with 80% ethanol and elution in 20 µL of nuclease-free water.

For second-strand synthesis, the reverse-transcribed sample was mixed with 25 µL of 2x LongAmp Taq Master Mix (NEB, M0533), 2 µL of 10 µM T7-ext primer, and 3 µL of nuclease-free water. The mixture was incubated under the following conditions: 94°C for 1 minute, 60°C for 1 minute, 65°C for 15 minutes, and held at 4°C. Clean-up was performed using a 0.8x AMPure XP bead ratio, followed by two washes with 80% ethanol and elution in 21 µL of nuclease-free water.

IVT was carried out on the generated cDNA using the New England Biolabs T7 High Yield RNA Synthesis Kit (NEB, E2040S) and their IVT protocol, without modification, on 14 µL of the cDNA. In-vitro transcribed mRNA was cleaned up using the New England Biolabs Monarch® Spin RNA Cleanup Kit (NEB, T2050). All resulting mRNA samples were sequenced using Nanopore direct RNA sequencing with the SQK-RNA004 kit on separate PromethION RNA flow cells.

### Models for m6A and pseU detection

#### Dorado

For Dorado, we directly fed the raw sequencing POD5 files to the Dorado basecaller (v0.7.2), using the rna004_130bps_sup@v5.0.0 model (v1, 4kHz), which was the latest model available at the time of evaluation, to detect both m6A and pseU modifications during the basecalling process. The output was a modBAM file, containing specific modification tags (MM and ML) that annotate the m6A/pseU positions and their corresponding probabilities. The modBAM file was aligned to the GENCODE(v46) primary assembly reference using Minimap2. Then we ran ModKit to extract the modification information from the aligned modBAM. The final output was a BED-like table that recorded the probability and sequence information of every predicted modified base on each read. Notably, we used ModKit solely to determine the recommended probability for each sample and did not rely on its comprehensive filtering strategy for per-site aggregation. Instead, we manually calculated per-site modification ratios based on selected thresholds.

#### m6Anet

For m6Anet, we first completed the basecalling of the raw POD5 files with the regular high-accuracy model from Dorado (without modification detection) and aligned the output to GENCODE comprehensive transcriptome reference (m6Anet requires BAM aligned to transcriptome) using Minimap2. We then followed the instructions from the latest release (v-2.1.0) of m6Anet. We preprocessed the data by converting and merging the raw POD5 files into BLOW5 format using f5c. The BLOW5 files, along with the reference and aligned BAM files, were used for f5c eventalign with rna004.nucleotide.5mer.model. This allowed us to proceed with m6Anet’s data preparation and inference using their latest model. The output consisted of two files: one BED-like table containing predictions for each base on every read, and another table providing the overall probability and modification fraction for each transcriptomic site based on their multi-instance learning algorithm.

However, since m6Anet runs on a transcriptome reference, its output positions are expressed in transcript coordinates. To compare these predictions with those from Dorado, which are in genomic coordinates, we performed an additional step to transform m6Anet’s transcript coordinates into genomic coordinates. Biologically, this procedure poses a challenge because m6Anet’s predictions, based on the transcriptome reference, do not include other genomic regions—particularly introns—whereas Dorado’s predictions, using a genomic reference, may encompass such regions. To ensure a fair comparison and maintain consistency in the regions considered, we limited the predicted sites to exonic regions and those with a DRACH motif, as m6Anet only considers these.

### Quality control and quantification of gene isoform expression

The raw sequencing BAM file after basecalling was processed by LongReadSum[36], which is a fast and flexible quality control and signal summarization tool for long-read sequencing data. For the quantification of gene isoform expression from our long-read sequencing output, we used LIQA[37] to calculate the corrected count of read per gene and rescaled the result by a TPM-like method for better visualization.

### Performance evaluation

As described in the results, we evaluated the performance of our modification detection tools (Dorado, m6Anet) using four primary metrics—recall, correlation, false positive rate (FPR), and false discovery rate (FDR). FPR and FDR are also calculated at both the per-read and per-site level. Below, we describe our assumptions and the precise definitions of these metrics.

#### Predictions and Ground Truth

We obtain per-read predictions directly from each model’s output. To derive per-site results, we apply per-read modification probability thresholds (see below) and aggregate the resulting calls for each genomic site. In our IVT control samples, all adenosines are assumed to be unmodified (true negatives). In WT samples, we rely on chemically validated benchmark datasets to identify truly modified positions, but we do not use these data to define true negatives because of potential noise and sensitivity limitations in the experimental methods. Finally, to ensure consistent comparisons, we align all per-site results (both predictions and ground truth) to the same genomic list of exonic DRACH sites with ≥10 ONT reads of coverage.

#### Thresholding

Each model outputs a per-read modification probability for each base. If that probability exceeds a chosen threshold (e.g., the model’s recommended cutoff), the base in that read is classified as “modified”; otherwise, it is deemed “unmodified.” By aggregating all read-level classifications that at the same genomic site, we obtain the “per-site modification ratio,” which then determines whether that site is ultimately classified as modified or unmodified. Sweeping the per-read modification threshold from 0 to 1 thus generates a range of recall, correlation, FPR, and FDR values.

#### Recall (sensitivity)

Recall measures the proportion of genuinely modified sites—according to the ground truth—that are correctly labeled as modified in the predictions. Formally,

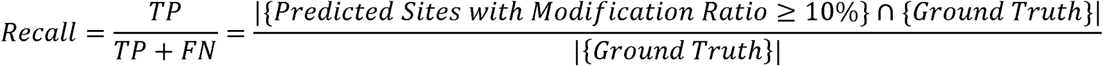

where TP (True Positives) are truly modified sites that the model correctly classifies as modified, and FN (False Negatives) are truly modified sites that the model fails to identify.

#### Correlation (i.e., Spearman r)

Correlation assesses how closely the predicted per site modification ratio aligns with a reference ratio (e.g., ground truth or another model’s prediction). All Spearman correlation coefficients are calculated on the common sites between groups.

#### False Positive Rate (FPR)

FPR quantifies the fraction of truly unmodified bases or sites (in the IVT samples) that are incorrectly classified as modified:

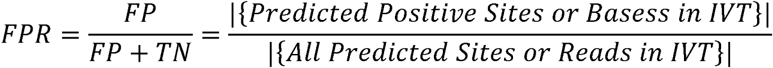

where FP (False Positives) are unmodified sites/bases from the IVT samples that the model mislabels as modified, and TN (True Negatives) are unmodified sites/bases correctly identified as unmodified. Per-Read FPR considers each unmodified bases per read (in IVT samples) that exceeds the threshold as an FP. Per-Site FPR considers each unmodified site (in IVT samples) for which the aggregated per site modification ratio is above a fixed cutoff (10%) as an FP.

#### False Discovery Rate (FDR)

Conventionally, FDR measures what fraction of all predicted positives are false. However, in our analysis, we cannot directly quantify this fraction; instead, we approximate it using the predicted positives observed in the IVT samples. This approximation follows a strategy discussed by Kong et al. [34], where IVT data serves as a practical surrogate for a fully negative dataset in scenarios where a complete gold standard is not available. Considering the difference between sample size, the positive counts are first normalized to fractions by dividing by the total counts of candidate sites:

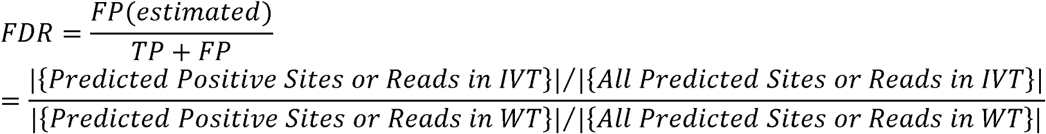

Per-Read FDR compares the fraction of predicted positive bases (i.e., bases classified as modified) between IVT and WT at the read level, effectively estimating how many of those “modified” calls in WT can be considered false based on IVT results. Per-Site FDR compares the fraction of predicted modified adenosine sites in IVT versus WT at the site level, again treating IVT signals as putative false calls.

## Competing Interests

The authors declare no competing interests.

## Availability of Data

The direct mRNA sequencing data has been deposited into ENA database under accession PRJEB80229.

## Author Contributions

ZZ and MUA performed the computational analysis. JC performed the sequencing and IVT experiments. MW prepared the samples, S-JG and YH provided samples and provided guidance on the study. KW supervised the execution of the study. ZZ drafted the initial version of the manuscript, and all authors revised the manuscript.

## Acknowledgements

We thank Wang lab members, especially Anagha Gouru, for valuable comments on the comparative evaluation project. This study is supported in part by NIH grant HG013359 (K.W.), CA279618 (Y.H.), and the CHOP Research Institute Omics Initiative (K.W.). We would like to thank the developers of GLORI and eTAM-seq in providing the results to be used in our comparative study on the same cell lines.

## Notes

### Competing Interest Statement

The authors have declared no competing interest.

