## Supplementary Materials for "A Comparative Evaluation of Computational Models for RNA modification detection using Nanopore sequencing with RNA004 Chemistry"

### Supplementary Material

Table 1. Summary of the direct RNA Nanopore sequencing data for wild-type (WT) and in-vitro transcribed (IVT) samples of HEK29T and HeLa cell lines.

| Cell Line | Group | Total Length (bp) | #Total Reads | #Mapped Reads | Mean/Median Mapped Read Length |
| --- | --- | --- | --- | --- | --- |
| HEK293T | WT | 9,018,509,026 | 8,613,942 | 5,680,018 | 1094.5/824 |
|  | IVT | 7,321,882,682 | 10,729,097 | 7,351,458 | 712.6/531 |
| HeLa | WT | 5,710,259,712 | 6,295,928 | 4,324,426 | 933.5/666 |
|  | IVT | 13,770,498,573 | 20,724,504 | 14,543,903 | 690.5/534 |

Table 2. Modkit estimated per-read modification probability threshold for each sample, used in Dorado predictions.

|  | HEK293T | HeLa |
| --- | --- | --- |
| WT | 0.607 | 0.619 |
| IVT | 0.627 | 0.623 |

Table 3. Performance metrics for RNA modification detection models (Dorado and m6Anet) and the size of ground truth datasets across wild-type HEK293T and HeLa cell lines. The table includes the total number of raw prediction output sites (#Total), predicted positive sites (#PP, under per site modification ratio >= 10%), true positive sites (#TP), false negative sites (#FN), as well as the recall for each model-cell line combination. Dorado generally shows higher F1 scores compared to m6Anet, reflecting better overall performance.

| Cell Line | Model/Dataset | #Total  (Raw Output) | #PP | #TP | #FN | recall |
| --- | --- | --- | --- | --- | --- | --- |
| HeLa | Dorado | 424,494 | 87,601 | 33,239 | 3,497 | 0.905 |
| HeLa | m6Anet | 204,831 | 72,839 | 18,663 | 18,073 | 0.508 |
| HEK293T | Dorado | 524,759 | 123,487 | 62,106 | 5,051 | 0.925 |
| HEK293T | m6Anet | 284,118 | 105,638 | 34,306 | 32,851 | 0.511 |
| HeLa | Ground Truth  (eTAM-seq) | 36736 |  |  |  |  |
| HEK293T | Ground Truth  (GLORI) | 67157 |  |  |  |  |

Figure 1. Scatter plots comparing isoform quantification in IVT versus WT samples for HEK293T (left) and HeLa (right). The Pearson correlation coefficients (r = 0.90 for HEK293T, r = 0.94 for HeLa) indicate that isoform abundances are highly consistent between IVT and WT samples in both cell lines.

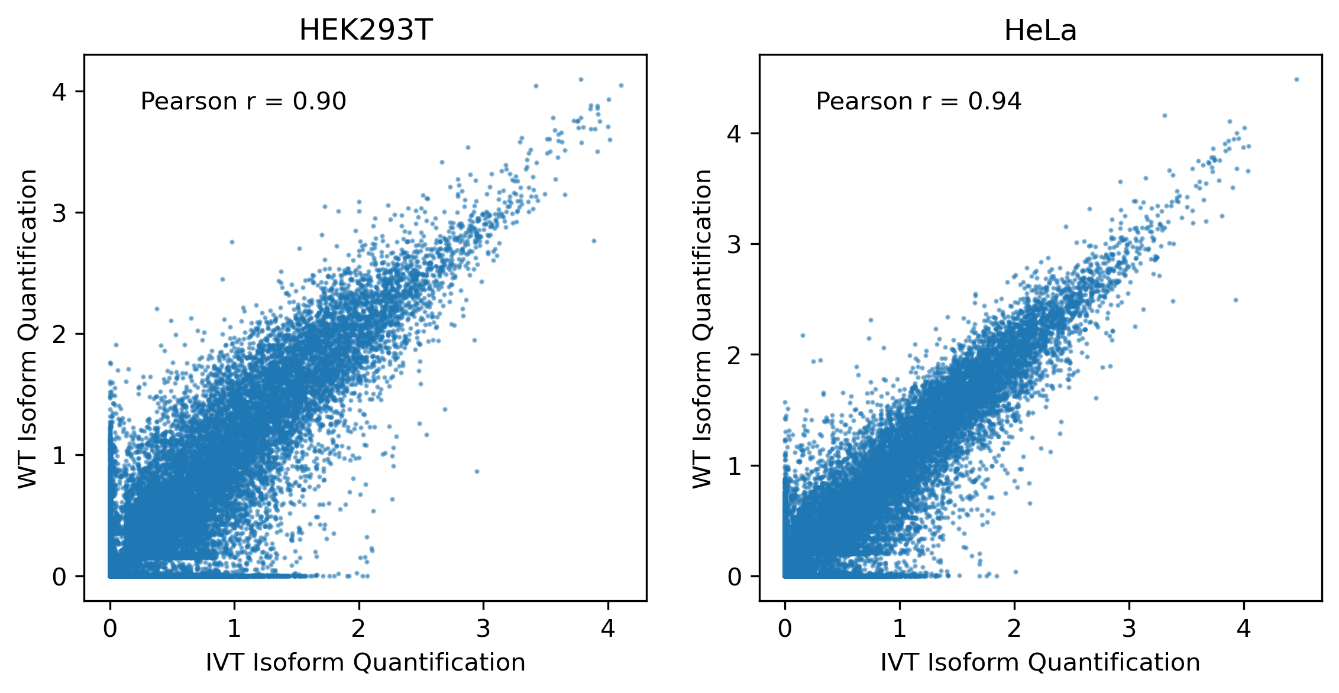

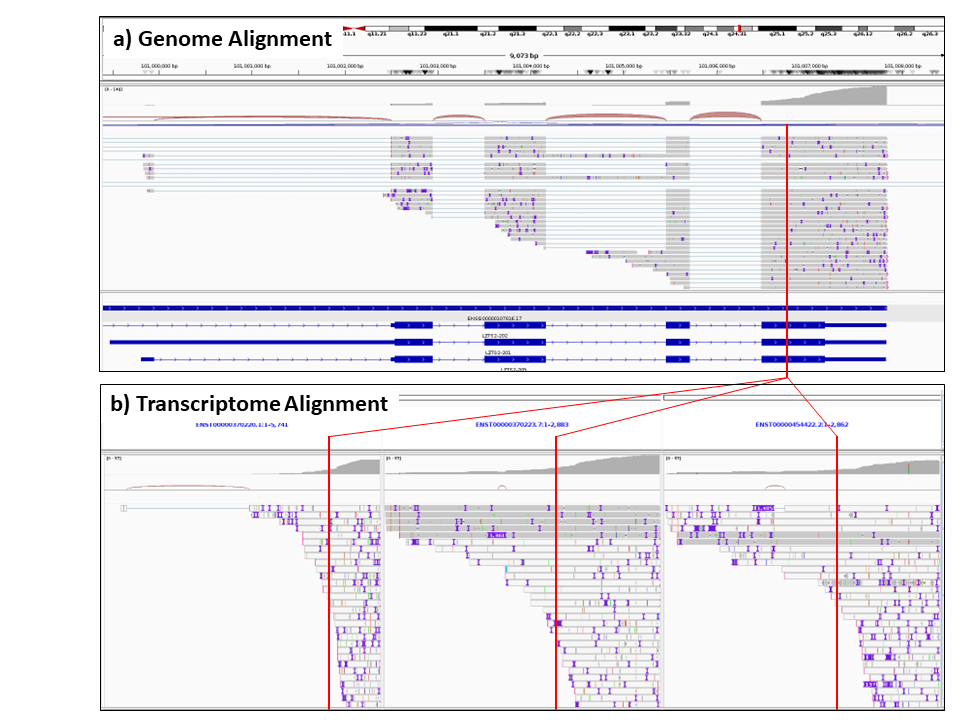
Figure 2.IGV plots of HEK293T wild-type ONT direct RNA reads aligned to LZST2 gene. a) shows reads aligned to GRCh38 reference genome sequence with splice-aware alignment. The vertical red line shows a DRACH site at chr10:101006765 and the genome annotation tracks in blue show three transcripts of LZST2. b) shows reads aligned to the spliced sequences of the three LZST2 transcripts, as required by m6Anet and f5c. The same DRACH site from reference genome is shown by vertical red lines in the three transcripts in b). Since most of the reads consist of the last 1-3 exons that are common between all transcripts, a majority of the read alignments in b) have mapping quality 0, shown by white highlighted horizontal reads, because these reads can be aligned equally as well to any of the transcripts. Whilst the reference DRACH motif has high coverage in a), each of the corresponding DRACH sites in b) have less than 20 coverage, which results in m6Anet not calling modification at any of these sites. For performance evaluation, we map any transcriptomic DRACH site modifications calls from m6Anet back to the genomic DRACH site, and merge predictions from all transcripts.

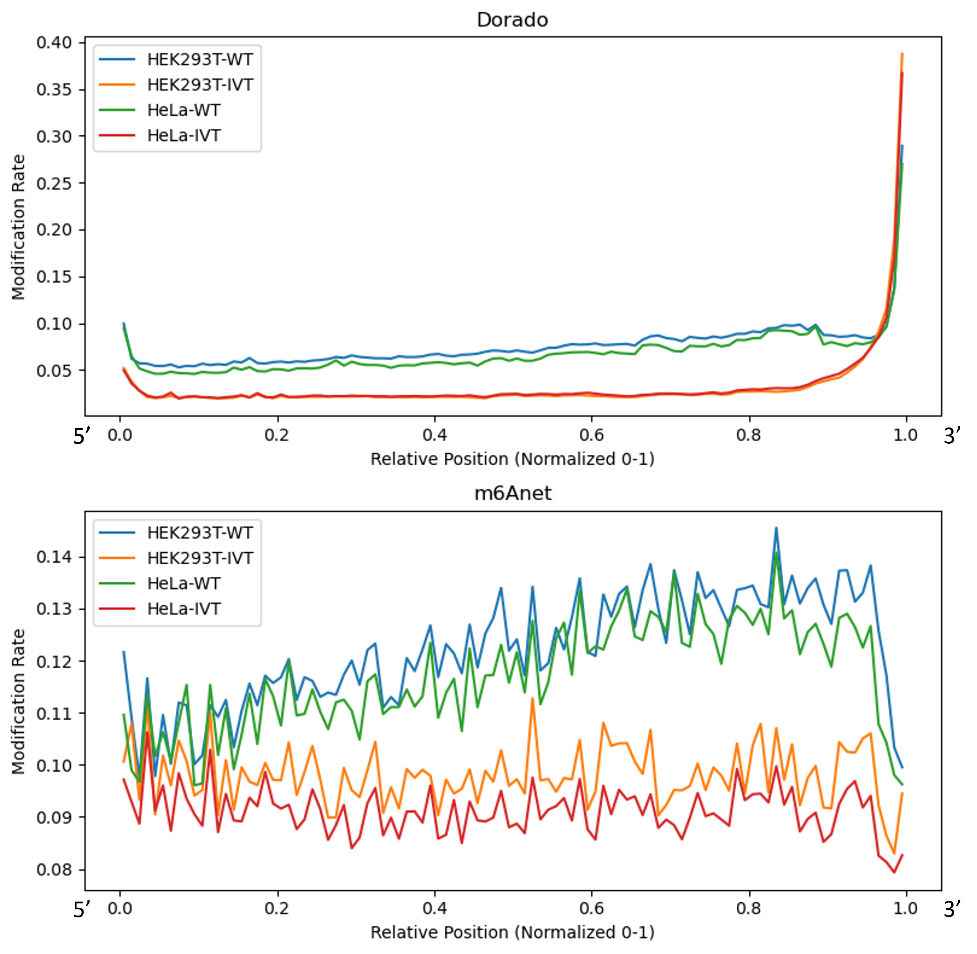

Figure 3. Predicted modification rate profiles along the normalized read length (from 5’ to 3’) for Dorado (top panel) and m6Anet (bottom panel). Each line represents either WT or IVT RNA from HEK293T or HeLa, color‐coded as shown in the legend. The x‐axis is normalized so that 0 and 1 correspond to the 5’ and 3’ ends of each read, respectively, while the y‐axis indicates the fraction of bases classified as modified at each position (represented by 100 bins between 0 and 1). Dorado’s predictions remain relatively low across most of the read but rise sharply at the 3’ end for both WT and IVT, whereas m6Anet shows a more gradual fluctuation throughout the read, with higher baseline levels in WT than IVT.

Figure 4. Density plots of inter-tool comparisons, correlating Dorado’s predicted modification ratios (x-axis) with m6Anet’s predicted modification ratios (y-axis) for HeLa (left) and HEK293T (right). The Spearman correlation coefficients and the number of sites is shown in each panel, highlighting broad agreement yet distinct differences at recommended probability thresholds.

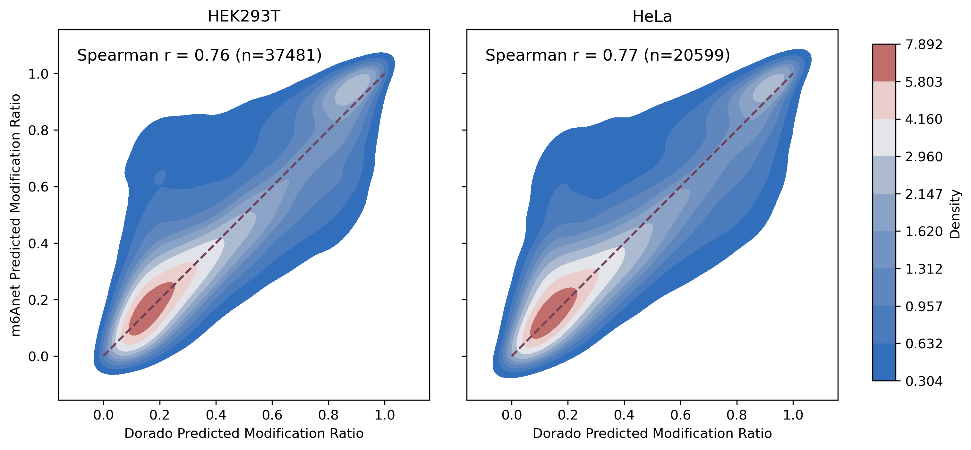
